# Diet composition of reintroduced Red-and-Green Macaws (*Ara chloropterus*) reflects gradual adaption to life in the wild

**DOI:** 10.1101/2021.04.14.439368

**Authors:** Noelia L. Volpe, Bettina Thalinger, Elisabet Vilacoba, Thomas W.A. Braukmann, Adrián S. Di Giacomo, Igor Berkunsky, Darío A. Lijtmaer, Dirk Steinke, Cecilia Kopuchian

## Abstract

Over the last two centuries, the Red-and-green Macaw (*Ara chloropterus*) has become locally extinct in Argentina. In an attempt to restore its key ecosystem functions as both disperser and regulator of large-seeded plants, a reintroduction project was initiated at the Iberá National Park in northeastern Argentina. The ability of released individuals to find food is crucial, in particular when working with captive-bred animals, as long-term establishment of a self-sustaining population depends on their short-term ability to exploit wild food sources. Monitoring of feeding habits is usually conducted through behavioral observation, but in recent years DNA metabarcoding has emerged as an alternative for obtaining highly resolved data on diet composition. In this study we use a combination of both techniques to characterize the breadth and composition of the reintroduced macaws’ diet. In addition, we evaluate the efficiency of both direct field observations and metabarcoding of feces as techniques to assess diet composition. Individuals fed on a variety of plant species (*n* = 49) belonging to a broad phylogenetic spectrum (28 families). Dietary richness estimated by direct observation and metabarcoding was similar, though smaller than the combination of the two datasets as both techniques detected at least 15 species not recorded by the other method. While the total number of detected species was higher for observational data, the rate of species-detection per sampling day was higher for metabarcoding. These results suggest that a combination of both methods is required in order to obtain the most accurate account of the total diversity of the diet of a species. The ability of the reintroduced macaws to successfully exploit local food resources throughout the year indicates a good level of adjustment to the release site, an important step towards the creation of a stable, self-sustaining population of Red-and-green Macaws in Northern Argentina.

**RESUMEN:** En el transcurso de los últimos dos siglos, el Guacamayo Rojo (*Ara chloropterus*) se ha extinguido en la Argentina. Buscando recuperar su rol ecológico tanto de dispersor como de depredador de semillas de gran tamaño, se comenzó un proyecto de reintroducción de la especie en el Parque Nacional Iberá, en la región noreste del país. La capacidad para encontrar alimento por parte de los individuos liberados es crucial, particularmente cuando se trabaja con animales provenientes de condiciones de cautiverio, ya que el establecimiento de una población autosuficiente a largo plazo dependerá de la habilidad de éstos para explotar fuentes de alimento silvestre a corto plazo. El monitoreo de hábitos alimenticios se realiza usualmente a través de observaciones comportamentales. Sin embargo, en los últimos años la técnica del meta-código de barras de ADN ha surgido como una alternativa para la obtención de datos de composición dietaria con alto nivel de resolución. En este estudio, utilizamos una combinación de ambas técnicas para caracterizar la amplitud y composición de la dieta de los guacamayos reintroducidos. A su vez, evaluamos la eficiencia de la observación directa y el código de barras genético de heces como técnicas para evaluar la composición de la dieta. Los individuos se alimentaron de una amplia variedad de especies (*n* = 49), abarcando un amplio espectro filogenético (28 familias). La riqueza dietaria estimada por observación directa y por meta-código de barras genético fue similar, aunque menor a la resultante de la combinación de todos los datos ya que ambas técnicas detectaron al menos 15 especies no registradas por el otro método. Mientras que el número total de especies detectadas fue mayor para los métodos observacionales, la tasa de detección de especies por día de muestreo fue mayor para el análisis genético. Estos resultados sugieren que una combinación de ambos métodos es necesaria para obtener la descripción más precisa posible de la diversidad dietaria total de una especie. La capacidad de los guacamayos reintroducidos para explotar recursos alimenticios locales a lo largo del año estaría indicando un buen nivel de adaptación al sitio de liberación, un paso muy importante hacia el establecimiento de una población de Guacamayo Rojo estable y autosuficiente en el norte de Argentina.

**Palabras clave:** *Ara chloropterus*, Conservación, Dieta, Frugivoría, Meta-código de barras, Guacamayo Rojo, Reintroducción, Ecología trófica

**LAY SUMMARY:**

- The Red-and-green Macaw reintroduction project aims to restore this species to Argentina, where it is locally extinct. To assess if reintroduced macaws are successfully adapting to life in the wild, we studied their foraging habits at the Iberá National Park. Their food consumption was observed visually, and their feces were analyzed with molecular methods for traces of the consumed plants.
- Macaws fed from a large diversity of food items, exhibiting a flexible diet which varied with fruit availability in different months. A combination of both methods was required to obtain the most accurate account of the total diversity of the diet of a species.
- The reintroduced macaws were able to successfully locate and exploit food resources throughout the year, indicating a good level of adjustment to the release site.

## INTRODUCTION

Over the last two centuries, Northern Argentina has experienced substantial defaunation mainly affecting large birds and mammals (Zamboni et al. 2017). One of the species that disappeared was the Red-and-green Macaw (*Ara chloropterus*), one of the largest species of the order Psittaciformes, last seen in the region almost 100 years ago and currently considered locally extinct in Argentina (Collar et al. 2020). Psittacids have traditionally been considered plant antagonists, acting as pre-dispersal predators by destroying seeds or removing them from the parent plants before they become viable (Trivedi et al. 2004, de Faria 2007, Ragusa-Netto 2011). Yet, an increasing body of literature highlights their importance as seed dispersers, being able to transport fruits across longer distances than smaller frugivores (Tella et al. 2015, Blanco et al. 2018). Thus, with the loss of the Red-and-green Macaw from the north of Argentina, its key roles as disperser and regulator of large-seeded plants in forests and savannas were removed from the region. In an attempt to restore these ecosystem functions, in 2014 the NGO Rewilding Argentina started the Red-and-green Macaw reintroduction project in the forests of the Iberá Wetlands, located in the north of the province of Corrientes (Zamboni et al. 2017).

One key consideration for the establishment of a new population in a reintroduction project is the ability of the animals to find food. Malnourished individuals will not only have a reduced chance of survival but also low reproductive success (Williams et al. 2013, Yu et al. 2015, Renton et al. 2015). Long-term establishment of a self-sustaining population depends on the short-term ability of released individuals to exploit wild food sources. For species that rely on seasonal resources, such as frugivores, this will involve not only locating food sources in space but also adapting to temporal changes in availability. Such dietary flexibility is a common characteristic of psittacid feeding ecology and is demonstrated by their broad diets (Renton et al. 2015). For the Red-and-green Macaw, this can mean relying on as many as 54 different plant species (Lee et al. 2014).

The Red-and-green Macaw reintroduction project relies entirely on captive-bred individuals which are naïve to foraging in the wild and will thus face particular challenges having to both find novel food sources and track their changes in availability along the year (Peignot et al. 2008). In social or gregarious species such as parrots, uptake of new food items by released individuals may be more successful if there are conspecifics already present in the area (Jones and Duffy 1993, Ewen et al. 2012). Unfortunately, the Iberá National Park currently lacks any native macaws or other large parrots from which reintroduced birds could learn to recognize food items. To compensate for this, we developed a pre-release training program to encourage captive-bred macaws to use wild fruits and seeds. Over the course of this program we were able to identify over 30 local plant species the macaws were willing to consume (N.L.V., personal observation). Although this indicates a potentially high diversity in the diet, it is likely that not all food items consumed within the captive environment will be eaten in the wild (Plair et al. 2013, Amaya-Villarreal et al. 2015). The realized dietary breadth for the free roaming population is determined by the ability of individuals to actually locate fruit bearing plants when they are available. Hence, only monitoring of released populations can help to evaluate if released macaws are succeeding at gathering food resources and to understand what key plant species are needed for the persistence of the macaw population in the area.

Monitoring of feeding habits is usually conducted through behavioral observation. Despite its common use, this technique comes with challenges and limitations because it is not always feasible to adequately track individuals, in particular when working with animals as mobile as macaws (Valentini et al. 2009a). The collected information is potentially incomplete, as the observer might not have witnessed enough feeding events or not have been able to tell if the foraged plants were actually ingested. DNA-based analysis of fecal matter can provide much more detailed information on consumed food items while also removing the need to track individuals for long periods of time (Valentini et al. 2009b, Oehm et al. 2011). In this context, metabarcoding has emerged as a powerful tool for obtaining highly resolved data on diet composition, which will be a reliable indicator of how well macaws are adapting to life in the wild. Several studies have reported the use of metabarcoding to analyze the diet of different animals, such as mammals (Lopes et al. 2020) and birds (Rytkönen et al. 2019, McClenaghan et al. 2019). However, despite the use of DNA barcoding to analyze seed dispersion (Lavabre et al. 2016, Galimberti et al. 2016), none of these studies applied metabarcoding to study the diet of frugivorous bird species. We consider this approach a valuable method to shed light on this field.

The main objective of this study was to describe the diet of reintroduced macaws at the Iberá National Park. In particular, we: 1) characterize dietary breadth and composition, and 2) evaluate the efficiency of both direct field observations and metabarcoding as useful techniques to assess diet composition.

## METHODS

### Study Site

The study was conducted at the Iberá National Park and the Iberá Provincial Reserve and Provincial Park (27.8704°S, 56.8801°W, Figure 1) in the province of Corrientes, Argentina, a wetland area consisting of flooded grasslands and savannas surrounding subtropical forest patches of variable sizes (0.02 – 11 ha). The climate is subtropical humid, with mild winters and no pronounced dry season. The average monthly temperature ranges from 15°C in June/July to 28°C in January/February with an average annual precipitation of 1800 mm (Neiff and Poi de Neiff 2006).

**FIGURE 1.**
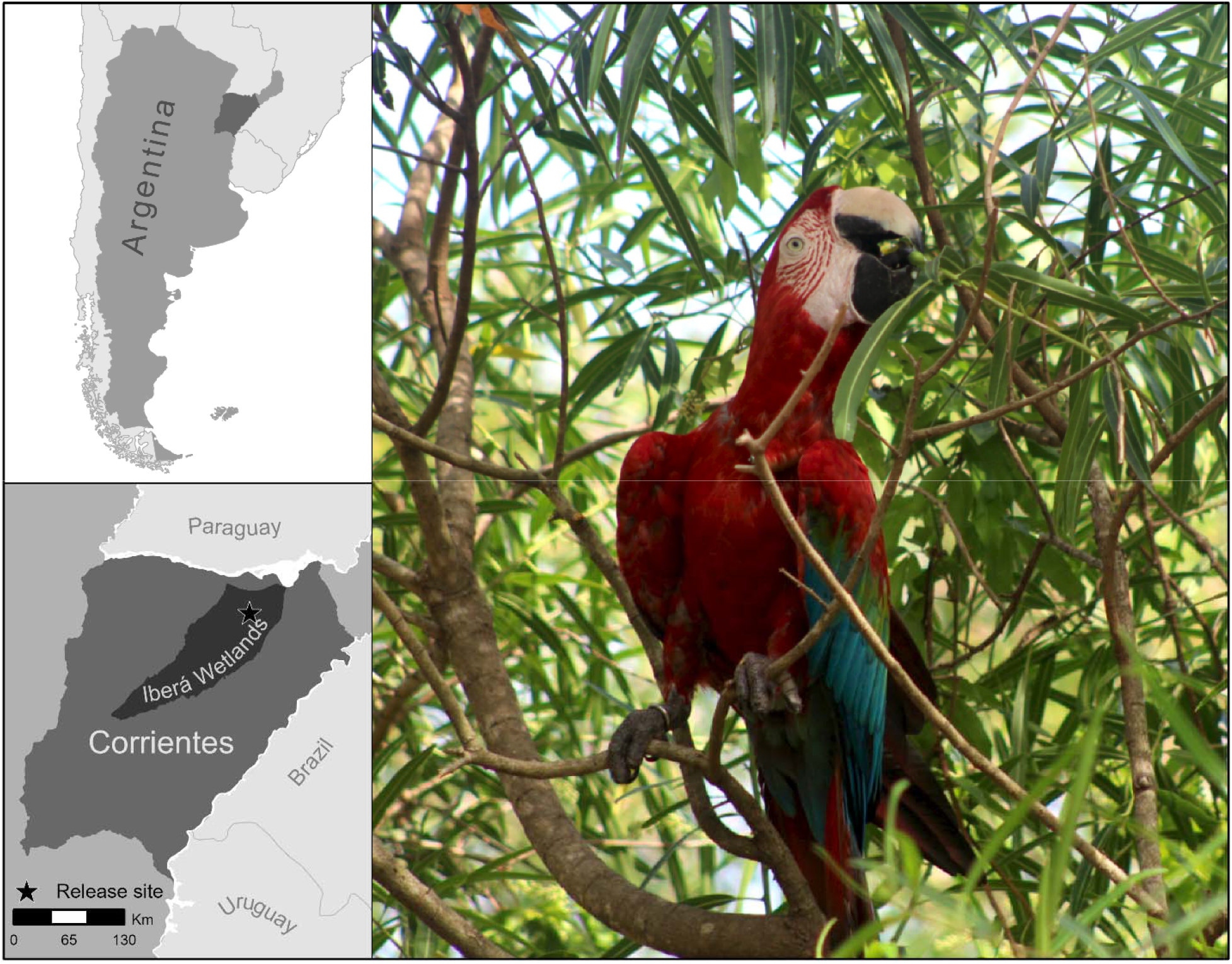
**Right**: Location of the study site, Portal Camby Retá (Iberá National Park), in the Iberá Wetlands region located in the province of Corrientes, northeastern Argentina. **Left**: reintroduced Red-and-green Macaw feeding on a wild fruit (*Sapium haematospermum*).

### Project Description

Macaws were donated to Rewilding Argentina by zoos, rescue centers and pet owners. They spent a quarantine period at the Aguará Conservation Centre (Corrientes, Argentina), where they were tested for a variety of diseases: mycoplasma, adenovirus, psittacine circovirus, Pacheco’s disease, paramyxovirus, influenza and chlamydia. In addition to this, individuals which showed signs of significant physical or behavioral problems (e.g., inability to fly, human imprinting) were removed from the candidate pool for release. Macaws deemed adequate were subsequently moved to a pre-release aviary at the release site in Portal Camby Retá (Iberá National Park), from where they were released after 11-16 months. For this study, we focused on two releases which took place in June 2017 (7 macaws; 4 females, 3 males) and in February 2018 (2 female macaws). However, visual feeding observations during pilot releases and DNA-based data retrieved from a sample collected in 2019 were included in the respective datasets. Thus, the time frame for observations and fecal sample collection was not identical. Food supplements consisting of a mixture of commercial fruits, vegetables and seeds provided on tree-platforms were available to the released macaws throughout the entire study period, applying a scheme of decreasing food supplementation over time: Four daily food supplements were offered until September 2017, three until December 2017, two until March 2018 and one from then onwards. This reduction in food supplementation was needed in order to motivate the macaws to expand their territory and forage from wild plants.

### Data Collection

#### Foraging observations

Between June 5 2017 and May 31 2018, we monitored the feeding activity of nine macaws, fitted with VHF radio-collars (Holohil AI-2C). Each macaw was followed using Yagi antennas for at least 4 consecutive hours each week for a total of 149 days. Every time we encountered macaws feeding on wild plants, we recorded date, time, GPS location, part eaten and species being consumed. We also included four additional foraging observations which took place during pilot releases in September 2015 and March 2017.

#### Fecal sampling and DNA barcode reference library

Feces were collected monthly between July 2017 and May 2018, with two additional samples collected in June 2019 (*n* = 10 macaws). Collections were made opportunistically while tracking the macaws with the aim to obtain at least one sample every two weeks, which led to a total of 96 samples collected on 61 different days (2 – 19 samples/month). Feces were collected between 1 and 40 minutes after defecation, except for two samples which were collected at a roost site early in the morning. Samples were stored at room temperature in 15 ml falcon tubes filled with ethanol (96 %) and after 1-3 months placed in a freezer at −20°C until further processing. Each tube contained from one (*n* = 86) to multiple (2 to 4; *n* = 10) fecal samples.

Based on observations of feeding preferences during the pre-release stage we built a DNA barcode reference library. We sampled 33 plant species that were expected to be used by macaws as food sources. Leaf plant tissue was taken from 3 individuals for all but three species for which we could only collect 1 or 2 samples. Samples were stored in silica gel until further processing.

### Sample processing

#### DNA barcoding and reference library compilation

The DNA extraction from plant tissues and amplification of barcode markers were performed at MACN following standard procedures; sequencing was done at the Centre for Biodiversity Genomics (CBG) at the University of Guelph in Canada. For a detailed description of these protocols see Supplementary Material 1A. In total, 65 of the 96 plant tissues produced ITS2 sequences corresponding to 24 different species; after filtering for contaminants and correcting for base-call errors, these were uploaded to BOLD (DS-IBERAFLO) and Genbank (accession numbers: MW845313-MW845377). Nine of the 33 sampled species expected to be eaten by the macaws failed to amplify. Sequences for these species, together with those of 179 other plant species occurring in the area (Arbo and Tressens 2002) were extracted from the ITS2 database hosted by the University of Würzburg (accessed 16th June 2020; Ankenbrand et al., 2015) in order to compile a custom reference database for maximum plausibility of the taxonomic assignment. In case sequences of specific species were not available, entries of the respective genus, family, tribe or order were used.

#### Metabarcoding of fecal samples

Metabarcoding of the fecal samples was carried out at the CBG and a detailed description of the entire process is contained in Supplementary Material 1B. Feces were processed in a laboratory dedicated to the handling of low-quality DNA samples with separate rooms for DNA extraction, PCR preparation and post-PCR processing. All DNA extracts were subjected to a metabarcoding approach using two consecutive PCRs and fusion primers (Elbrecht and Steinke 2019). PCR conditions were optimized for maximum yield of target length fragments, while minimizing the occurrence of non-target bands. The first round of PCR employed the primers ITS-u3 5’-CAWCGATGAAGAACGYAGC-3’ and ITS-u4 5’-RGTTTCTTTTCCTCCGCTTA-3’ (Cheng et al. 2016) and in the second PCR Illumina sequencing adapters were added using individually tagged fusion (Elbrecht and Steinke, 2019; Supplementary Material Table S1). Sequencing was carried out by the Advanced Analysis Centre at the University of Guelph using a 600 cycle Illumina MiSeq Reagent Kit v3 and 5% PhiX spike in. Sequencing results were uploaded to the Sequence Read Archive (SRA, Genbank, accession: PRJNA695029).

#### Bioinformatic analyses

Resulting sequence data were processed using the JAMP pipeline v0.67 (github.com/VascoElbrecht/JAMP). Sequences were demultiplexed, paired-end reads merged using Usearch v11.0.667 with fastq_pctid=75 (Edgar 2010), reads outside a 100 bp to 430 bp range were discarded and primer sequences trimmed by using Cutadapt v1.18 with default settings (Martin 2011). Sequences with poor quality were removed using an expected error value of 1.5 (Edgar and Flyvbjerg 2015) as implemented in Usearch. All sequences with less than five reads were removed during the denoising process. The obtained haplotypes were mapped against the custom sequence database; those without matches were subsequently blasted. Detailed information on the mapping process and determination of the levels of taxonomic resolution can be in Supplementary Material 1C.

The detected taxa were classified into five categories: Resource = wild local plants known to be a food item or considered likely to be so based on its characteristics (fruit-producing tree or vine); Provided = commercial fruits or vegetables included in the daily food supplements; Provided/Resource = level of resolution did not allow to exclude either option; Contamination = unlikely to have been eaten by the macaws (included algae and herbaceous or aquatic plants) and Ambiguous = could be either of the previous categories. Only taxa classified as Resource were included in the dietary analysis.

### Data Analysis

Diet breadth was estimated as the number of wild species consumed by the macaws, including a) species observed being eaten by the macaws, b) resource taxa detected in the feces resolved to the species level and c) resource taxa resolved to the genus level but not detected at the species level (e.g., *Tabebuia* sp. but not *Psidium* sp., as the latter is already represented by *Psidium guajava*). Given that pine trees in forestry plantations are the only conifers present in the study area, all conifer reads were treated as the same unit (*Pinus* sp.) regardless of their level of resolution (order, family, genus or species). We ranked the relative importance of each consumed species by estimating their proportion of occurrence in the diet, *i.e.* the number of days during which a given food item was detected over the total number of sampling days (feces-collection days and observation days; *n* = 153). To test whether macaws consumed each species according to their availability in the area, we used the Spearman rank correlation to evaluate the relationship between the number of feeding events and the proportion of fruiting trees of 14 of the consumed species for which phenological information was available (see Supplementary Material Figure S2). In order to compare results from both sampling methodologies, we estimated the detection rate for each technique and used Pearson’s correlations to assess if foraging observations and fecal sampling data collected on the same dates led to similar conclusions regarding changes in resource use patterns over time. All data were analyzed in R 3.6.2 (R Core Team 2020) using *tidyverse* and associated packages (Wickham et al. 2019). Results are expressed as mean ± standard error.

## RESULTS

### Foraging Observations

During the study period we recorded feeding bouts on 140 out of the 149 observation days, adding up to 336 hours of records on feeding behavior (*n =* 551 feeding events). Macaws fed on 536 different individual trees, as well as five vines and one epiphyte. Macaws consumed mainly fruits and seeds (98.5% of the events, 29 species), although they were occasionally observed chewing on flowers (0.9% of the events, four species used) and leaves (0.6% of the events, one species used).

### Fecal Analysis

Illumina sequencing produced 23.2 million paired-end reads. Of the 96 samples, 13 did not contain reads which passed the quality filtering process. Of the 8.1 million reads that passed all quality filtering and denoising steps, 1.8 million (22%) could be assigned to sequences in the custom reference database. The percentage of assignable reads varied between 0 and 99% (average 24%) for the individual samples. After blasting previously unmapped sequences, 3.1 million (38%) did not result in a clear taxonomic assignment, 2.4 million (30%) were assigned to fungi and 0.07 million (0.88%) matched to Viridiplantae. The majority of plant reads (custom database plus Genbank) were assigned to species (57%); 17%, 21% and 5%, to genus, family, and order respectively (Figure 2; see Supplementary Material Table S3 for information on individual samples). Of all negative controls only 4 (2 PCR controls, the fume hood (evaporation) control and the extraction control) contained reads which passed quality filtering. However, only 10 reads (of one PCR control) could be mapped (to *Lens culinaris*). As the read numbers in fecal samples assigned to *Lens culinaris* were always more than twice as high, we refrained from correcting the read numbers and occurrence data.

**FIGURE 2.**
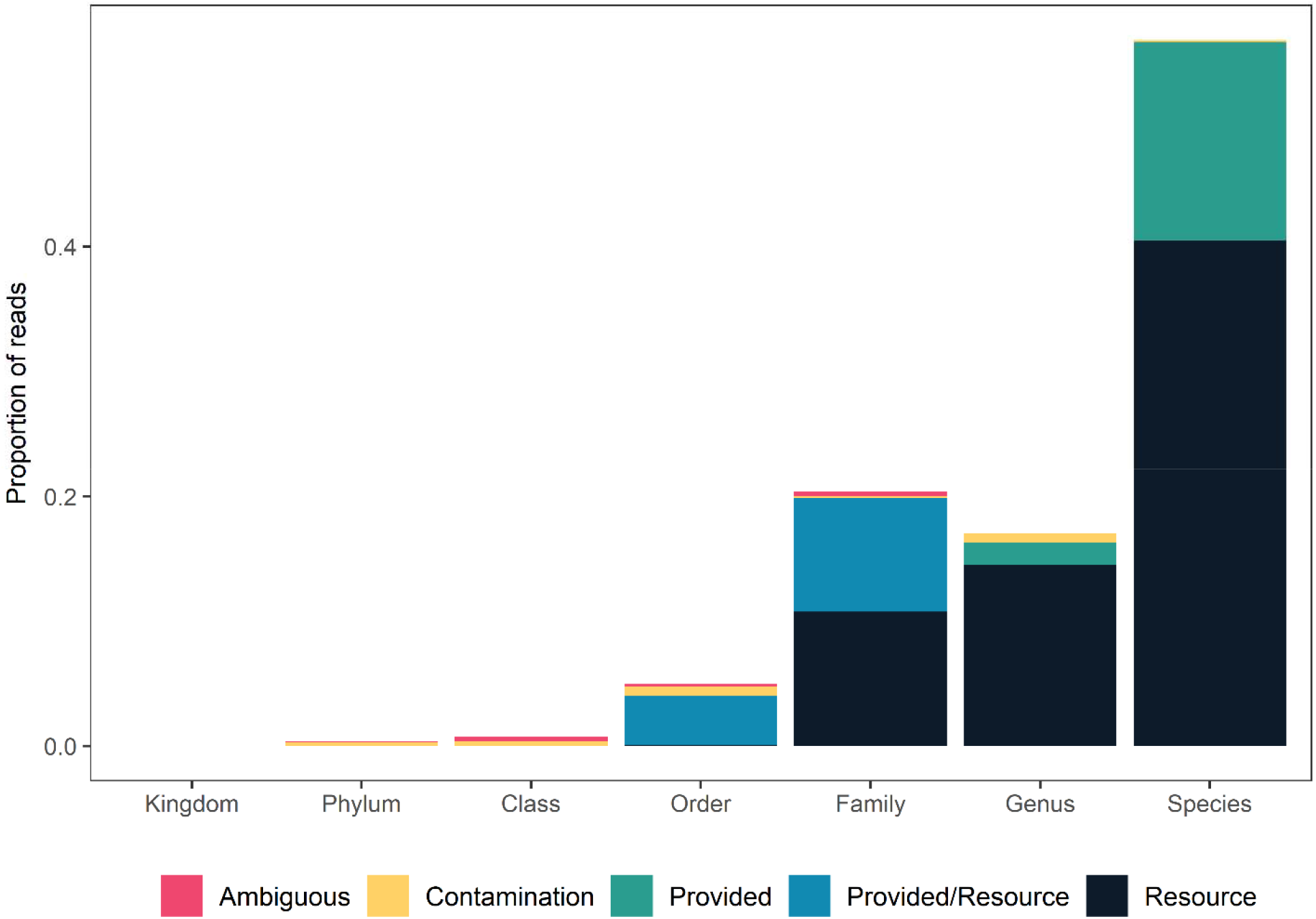
Distribution of reads among taxonomic levels and categories of use, based on 1,840,811 reads from 83 samples (**Resource** = wild local plants; **Provided** = commercial fruits or vegetables; **Provided/Resource** = level of resolution does not allow to exclude either option; **Contamination** = Accidental presence, byproduct of fecal sample collection; **Ambiguous** = either of the previous categories).

On average, 4.63 ± 0.29 species were detected per sample (*n* = 81, range = 1 – 10 species, Figure 3A). When taking into account just the resource taxa, the average dropped to 2.66 ± 0.16 species per sample (*n* = 75, range = 1 – 6 species, Figure 3B). The majority of reads corresponded to confirmed food resources, but provided food was detected in 72% of the samples (Figure 4A). Similarly, provided food was present in the feces throughout all sampling months although its relative presence decreased towards the end of the study period, coinciding with the reduction in food supplementation (Figure 4B).

**FIGURE 3.**
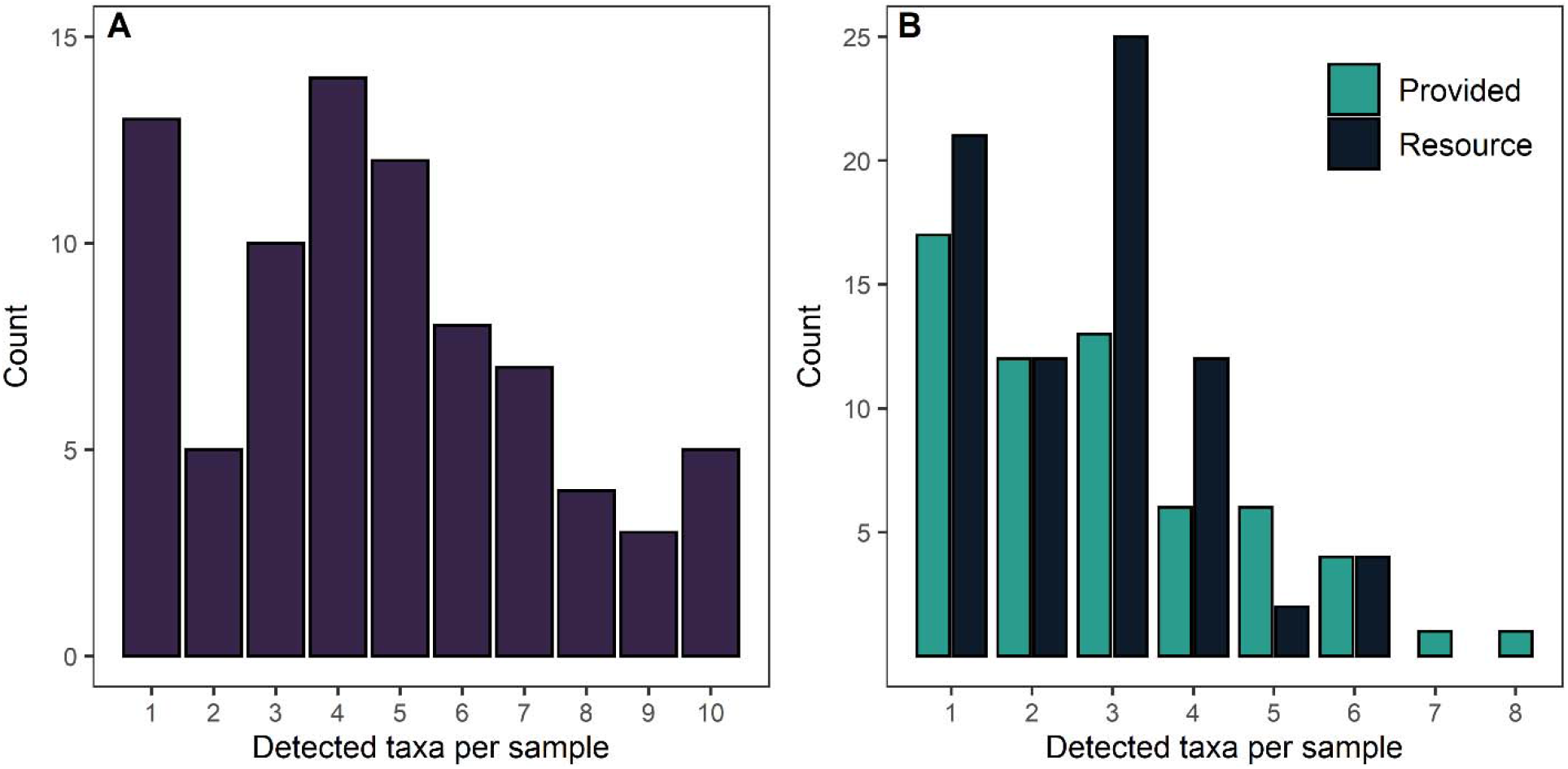
Plant diversity in Red-and-green Macaw feces collected at the Iberá Wetlands. (A) Distribution of the total number of taxa detected at the species or genus level per sample; (B) Distribution of the number of provided taxa and resource taxa detected at the species or genus level per sample (*n* = 83 samples).

**FIGURE 4.**
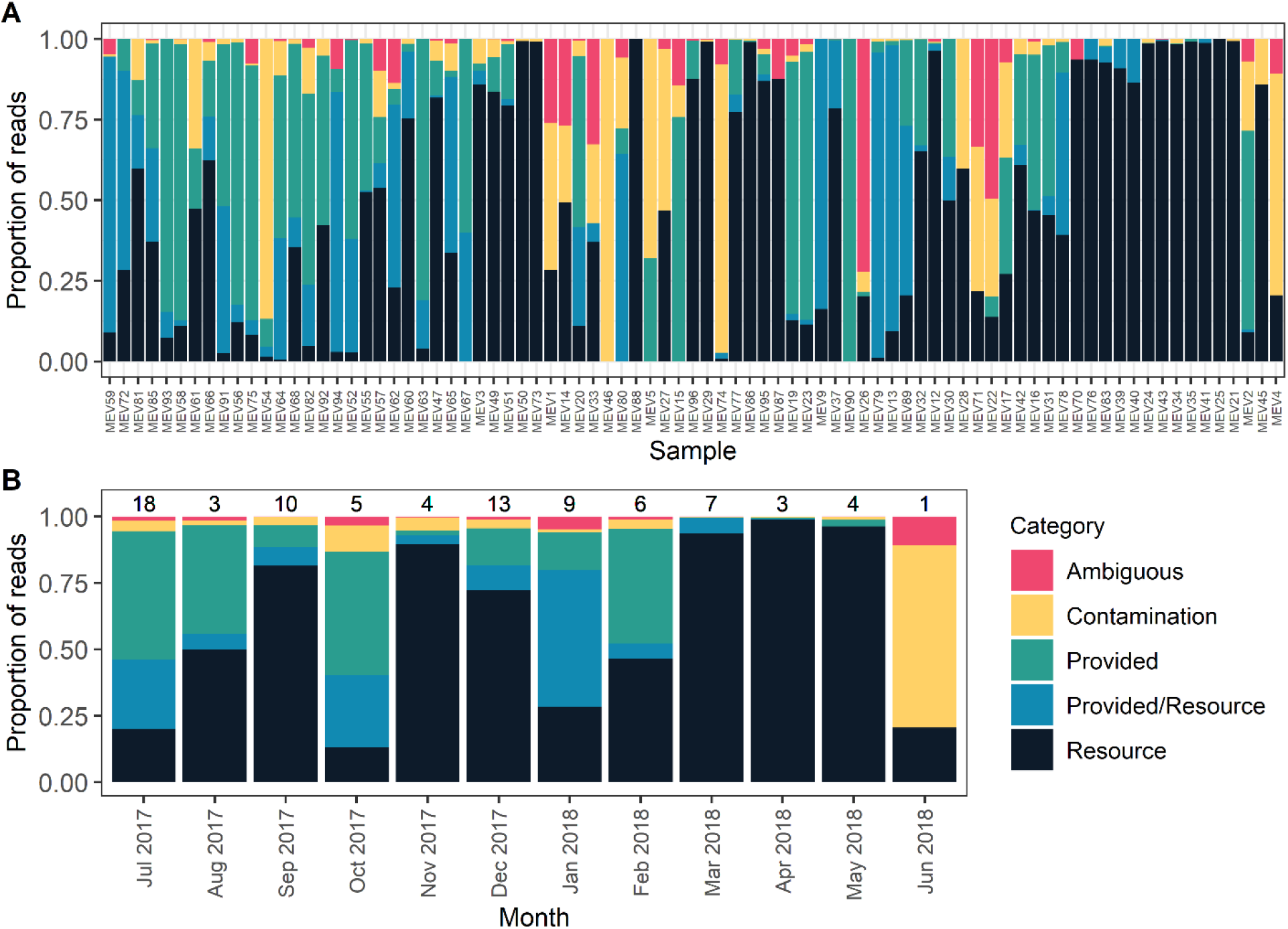
Distribution of plant reads proportions between samples (**A**) and months (**B**). Samples in the upper panel (**A**) are sorted based on their collection date (Jul-15-2017 to Jun-2-2018). Values above the columns in (**B**) represent the number of samples collected that month (*n* = 83 samples). (**Resource** = wild local plants; **Provided** = commercial fruits or vegetables; **Provided/Resource** = level of resolution does not allow to exclude either option; **Contamination** = Accidental presence, byproduct of fecal sample collection; **Ambiguous** = either of the previous categories).

### Diet Composition

Macaws exhibited a diverse diet feeding on 49 plant species from 28 different families (Table 1). Most of the feeding activity was concentrated on a small number of species, which were detected on more than 15 sampling dates. Particularly important was *Psidium guajava*, which was detected on 59% of the days. *Syagrus romanzoffiana* and *Inga edulis* were detected over 26% and 20% of the days, respectively, while *Ficus luschnathiana*, *Enterolobium contortisiliquum*, *Croton urucurana*, *Sapium haematospermum*, *Pinus* sp. and *Melia azedarach* appeared in the diet on 10-14% of the days. The 40 remaining species were present in the diet on less than 10% of the sampling days. The monthly recurrence of use of the different food items varied between species. While most of them were detected as being consumed only during one month, others were part of the diet for most of the year (e.g., *P. guajava*, *S. romanzoffia* and, *I. edulis*; Table 1).

**Table 1.** Species used by Red-and-green Macaws released at the Iberá National Park; **Part:** part of the plant consumed (only for direct observations); **Method:** technique used to determine presence in the diet (O = Observation, M = Metabarcoding); **PO:** proportion of occurrence (percentage of sampling days on which the species was detected in the diet); **Month:** month during which feeding was detected by direct observation (black), metabarcoding (light grey) or both methods (dark grey); **Total:** number of days sampled in a month. * Part: rs = ripe seed, us = unripe seed, rp = ripe pulp, up = unripe pulp r = ripe fruit (unclear if seed or pulp), u = unripe fruit (unclear if seed or pulp), f = flower, l = leaf, a = aril.

| Family | Scientific name | Part* | Method | PO | 2017 |  |  |  |  |  |  | 2018 |  |  |  |  |
| --- | --- | --- | --- | --- | --- | --- | --- | --- | --- | --- | --- | --- | --- | --- | --- | --- |
|  |  |  |  |  | J | J | A | S | O | N | D | J | F | M | A | M |
| Myrtaceae | *Psidium guajava* | up,rp,us,rs | O,M | 59 |  |  |  |  |  |  |  |  |  |  |  |  |
|  | *Eugenia myrcianthes* | up,rp,us,rs | O,M | 6 |  |  |  |  |  |  |  |  |  |  |  |  |
|  | *Eugenia uniflora* | rp | O,M | 5 |  |  |  |  |  |  |  |  |  |  |  |  |
|  | *Blepharocalyx salicifolius* | - | M | 1 |  |  |  |  |  |  |  |  |  |  |  |  |
|  | *Eugenia pitanga* | - | M | 1 |  |  |  |  |  |  |  |  |  |  |  |  |
| Fabaceae | *Inga edulis* | us,rs,a | O | 20 |  |  |  |  |  |  |  |  |  |  |  |  |
|  | *Enterolobium contortisiliquum* | rp,rs | O | 13 |  |  |  |  |  |  |  |  |  |  |  |  |
|  | *Delonix regia* | rs,f | O | 3 |  |  |  |  |  |  |  |  |  |  |  |  |
|  | *Chloroleucon tenuiflorum* | us | O | 1 |  |  |  |  |  |  |  |  |  |  |  |  |
|  | *Senna* sp. | - | M | 1 |  |  |  |  |  |  |  |  |  |  |  |  |
| Euphorbiaceae | *Croton urucurana* | us | O,M | 14 |  |  |  |  |  |  |  |  |  |  |  |  |
|  | *Sapium haematospermum* | us,rs | O,M | 12 |  |  |  |  |  |  |  |  |  |  |  |  |
|  | *Alchornea triplinervia* | us | O,M | 7 |  |  |  |  |  |  |  |  |  |  |  |  |
|  | *Sebastiania commersoniana* | us | O,M | 2 |  |  |  |  |  |  |  |  |  |  |  |  |
| Arecaeae | *Syagrus romanzoffiana* | up,rp,us,rs | O | 26 |  |  |  |  |  |  |  |  |  |  |  |  |
|  | *Acrocomia aculeata* | us,rs | O | 1 |  |  |  |  |  |  |  |  |  |  |  |  |
| Moraceae | *Ficus luschnathiana* | rp,rs | O,M | 14 |  |  |  |  |  |  |  |  |  |  |  |  |
| Pinaceae | *Pinus* sp. | us | O,M | 11 |  |  |  |  |  |  |  |  |  |  |  |  |
| Meliaceae | *Melia azedarach* | us,rs | O,M | 10 |  |  |  |  |  |  |  |  |  |  |  |  |
| Lauraceae | *Ocotea diospyrifolia* | rs | O | 8 |  |  |  |  |  |  |  |  |  |  |  |  |
| Bignoniaceae | *Jacaranda mimosifolia* | us | O | 3 |  |  |  |  |  |  |  |  |  |  |  |  |
|  | *Handroanthus impetiginosus* | f | O | 1 |  |  |  |  |  |  |  |  |  |  |  |  |
|  | *Tabebuia* sp. | - | M | 1 |  |  |  |  |  |  |  |  |  |  |  |  |
|  | *Dolichandra unguis-cati* | f | O | 1 |  |  |  |  |  |  |  |  |  |  |  |  |
| Solanaceae | *Cestrum nocturnum* | - | M | 3 |  |  |  |  |  |  |  |  |  |  |  |  |
|  | *Cestrum parqui* | u | O | 2 |  |  |  |  |  |  |  |  |  |  |  |  |
| Tiliaceae | *Luehea divaricata* | us | O | 3 |  |  |  |  |  |  |  |  |  |  |  |  |
| Sapotaceae | *Chrysophyllum gonocarpum* | u,r | O | 3 |  |  |  |  |  |  |  |  |  |  |  |  |
|  | *Pouteria gardneriana* | - | M | 1 |  |  |  |  |  |  |  |  |  |  |  |  |
| Rhamnaceae | *Sageretia elegans* | r | O,M | 3 |  |  |  |  |  |  |  |  |  |  |  |  |
|  | *Gouania polygama* | us,rs | O | 1 |  |  |  |  |  |  |  |  |  |  |  |  |
| Asteraceae | *Mikania cordifolia* | - | M | 3 |  |  |  |  |  |  |  |  |  |  |  |  |
| Viscaceae | *Phoradendron bathyoryctum* | r | O | 2 |  |  |  |  |  |  |  |  |  |  |  |  |
| Urticaceae | *Cecropia pachystachya* | l | O,M | 1 |  |  |  |  |  |  |  |  |  |  |  |  |
|  | *Urera* sp. | - | M | 1 |  |  |  |  |  |  |  |  |  |  |  |  |
| Sapindaceae | *Allophylus edulis* | - | M | 1 |  |  |  |  |  |  |  |  |  |  |  |  |
|  | *Mangifera indica* | - | M | 1 |  |  |  |  |  |  |  |  |  |  |  |  |
|  | *Paullinia elegans* | rs | O | 1 |  |  |  |  |  |  |  |  |  |  |  |  |
| Apocynaceae | *Tabernaemontana catharinensis* | - | M | 1 |  |  |  |  |  |  |  |  |  |  |  |  |
| Annonaceae | *Annona emarginata* | rp | O | 1 |  |  |  |  |  |  |  |  |  |  |  |  |
| Verbenaceae | *Vitex* sp. | - | M | 1 |  |  |  |  |  |  |  |  |  |  |  |  |
| Salicaceae | *Xylosma venosa* | r | O | 1 |  |  |  |  |  |  |  |  |  |  |  |  |
| Rubiaceae | *Psychotria carthagenensis* | - | M | 1 |  |  |  |  |  |  |  |  |  |  |  |  |
| Primulaceae | *Myrsine* sp. | - | M | 1 |  |  |  |  |  |  |  |  |  |  |  |  |
| Phytolaccaceae | *Phytolacca dioica* | - | M | 1 |  |  |  |  |  |  |  |  |  |  |  |  |
| Passifloraceae | *Passiflora caerulea* | rp,rs,f | O | 1 |  |  |  |  |  |  |  |  |  |  |  |  |
| Melastomataceae | *Miconia pusilliflora* | r | O,M | 1 |  |  |  |  |  |  |  |  |  |  |  |  |
| Malvaceae | *Theobroma* sp. | - | M | 1 |  |  |  |  |  |  |  |  |  |  |  |  |
| Cucurbitaceae | *Lagenaria* sp. | - | M | 1 |  |  |  |  |  |  |  |  |  |  |  |  |
| Total |  |  |  |  | 9 | 20 | 9 | 16 | 14 | 13 | 19 | 12 | 10 | 11 | 7 | 9 |

The relationship between resource use and availability varied between species. Plants with relatively short fruiting periods such as *Eugenia myrcianthes* and members of the family Euphorbiaceae, or those that were eaten mainly at their ripe stage, such as *Ocotea diospyrifolia*, *F. luschnathiana*, and *I. edulis* were consumed as they became available, with the number of feeding events correlated to the monthly availability of each species (Spearman rho = 0.63 – 0.83, *P*<0.05). On the other hand, the intensity of use of species with longer fruiting periods was less predictable and not associated to their availability (Spearman rho = − 0.02 – 0.56, *P*>0.05). Some species, like *P. guajava*, were used in high proportions even in months during which they had a low relative abundance, while others were only used for a few months (*Enterolobium contortisiliquum*) despite being available for most part of the year (Supplementary material Table S2).

### Comparison Between Techniques

Of the 49 plant species identified as being part of the macaw diet, 13 were detected by both techniques, while 17 were detected only by metabarcoding (7 of them resolved to genus) and 19 only by direct observation (Table 1, Fig 4). Of these 19 species not detected by metabarcoding, 7 could actually not be detected because they were not present in the database used for species identification. Both techniques identified *P. guajava* as the most important species for the macaws, but the relative use of the remaining shared species was more variable. For example, *Croton urucurana* was consumed in only 4% of the sampling days based on the observational data, but on 25% of the days based on the metabarcoding results (Figure 5).

**FIGURE 5:**
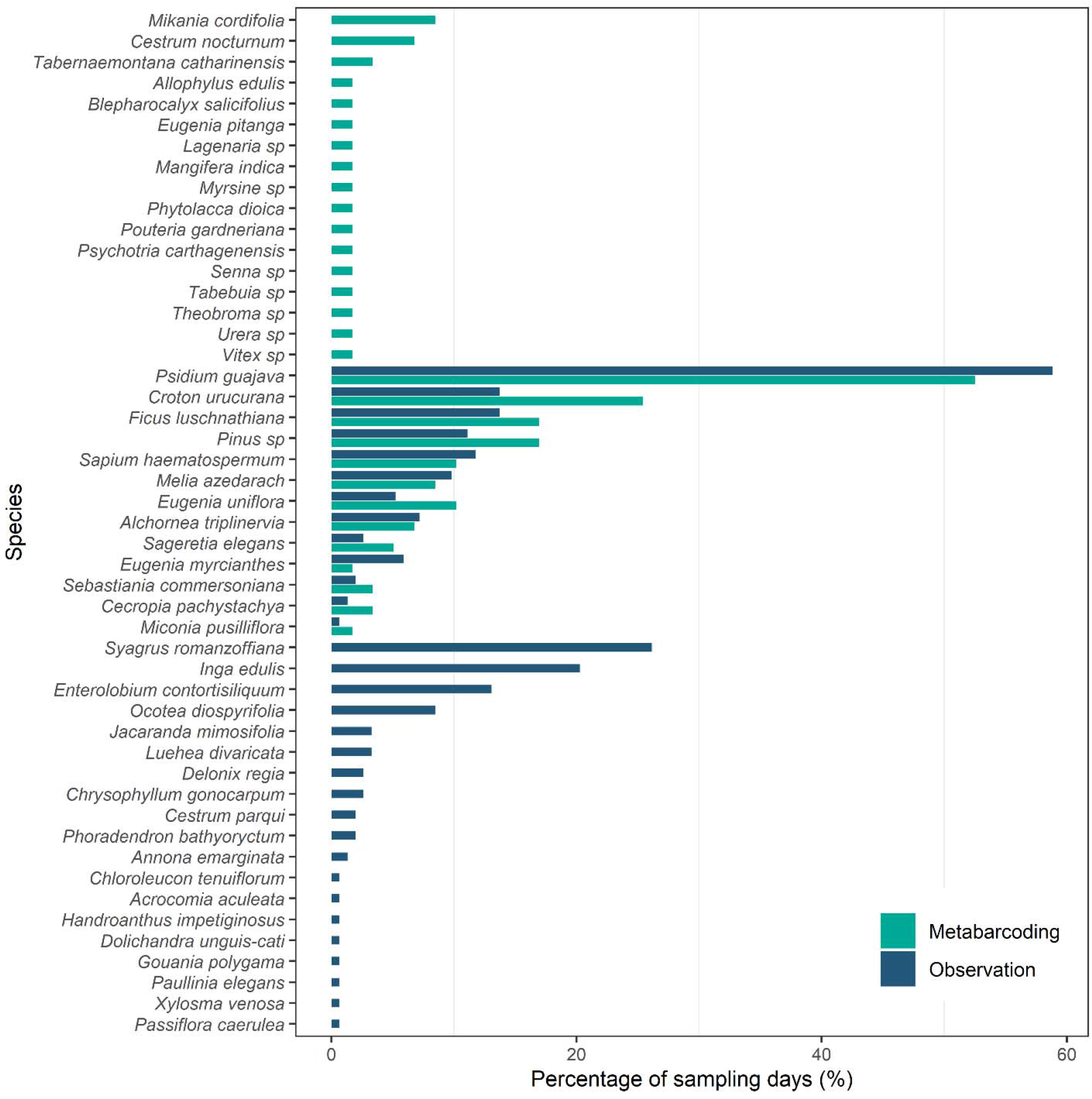
Plant species consumed by the Red-and-green Macaw at the Iberá Wetlands and their relative contribution to the diet (percentage of sampling days in which each species was detected) based on observational (blue; *n* = 140 days) and DNA-based data (teal; *n* = 53 days).

While the total number of detected species was higher for observational data, the rate of species-detection per sampling day was higher for metabarcoding. When comparing data collected on the same dates (*n* = 39 days) metabarcoding detected 28 species (0.72 species day-1) while observational data detected 22 species (0.56 species day-1). The difference between techniques was more pronounced when looking at detection rates each month, with an average of 1.81 ± 0.27 species detected per feces collection day (*n* = 11; range = 1 – 4 species), compared to only 1.35 ± 0.11 species per observation day (*n* = 11, range = 0.8 – 2 species).

Results from both techniques exhibited a similar pattern regarding the rate of increase in dietary richness since the date of release (Pearson correlation, *n* = 39, *r* = 0.95, *P*<0.05), showing a sharp initial growth during the first months and a flattening of the curve at around the 7-month mark, both leading to a similar final diet breadth estimate of 30-32 species (Figure 6). The overall pattern of changes in dietary breadth along the year was also similar between techniques (Pearson correlation, *n* = 11, *r* = 0.6, *P*<0.05), with peaks in July, September and December, followed by a decrease towards Autumn (Figure 7). The pattern of use of specific food items along the year differed between techniques, with observational data underestimating the use of many of them. At least ten species occurred in the diet of the macaws for much longer than observed (Figure 8). Metabarcoding revealed that the group of macaws began to eat some of the species several months before we first observed them doing so (e.g., *C. urucurana, Pinus* sp.) or did it for 1-7 months longer than recorded (e.g., *Eugenia uniflora, Ficus luschnathiana*). In the case of *Cecropia pachystachya*, its use was only observed during the pilot release of 2015, but metabarcoding highlighted that the macaws were also eating it during the 2017-2018 releases, though it was never visually detected.

**FIGURE 6.**
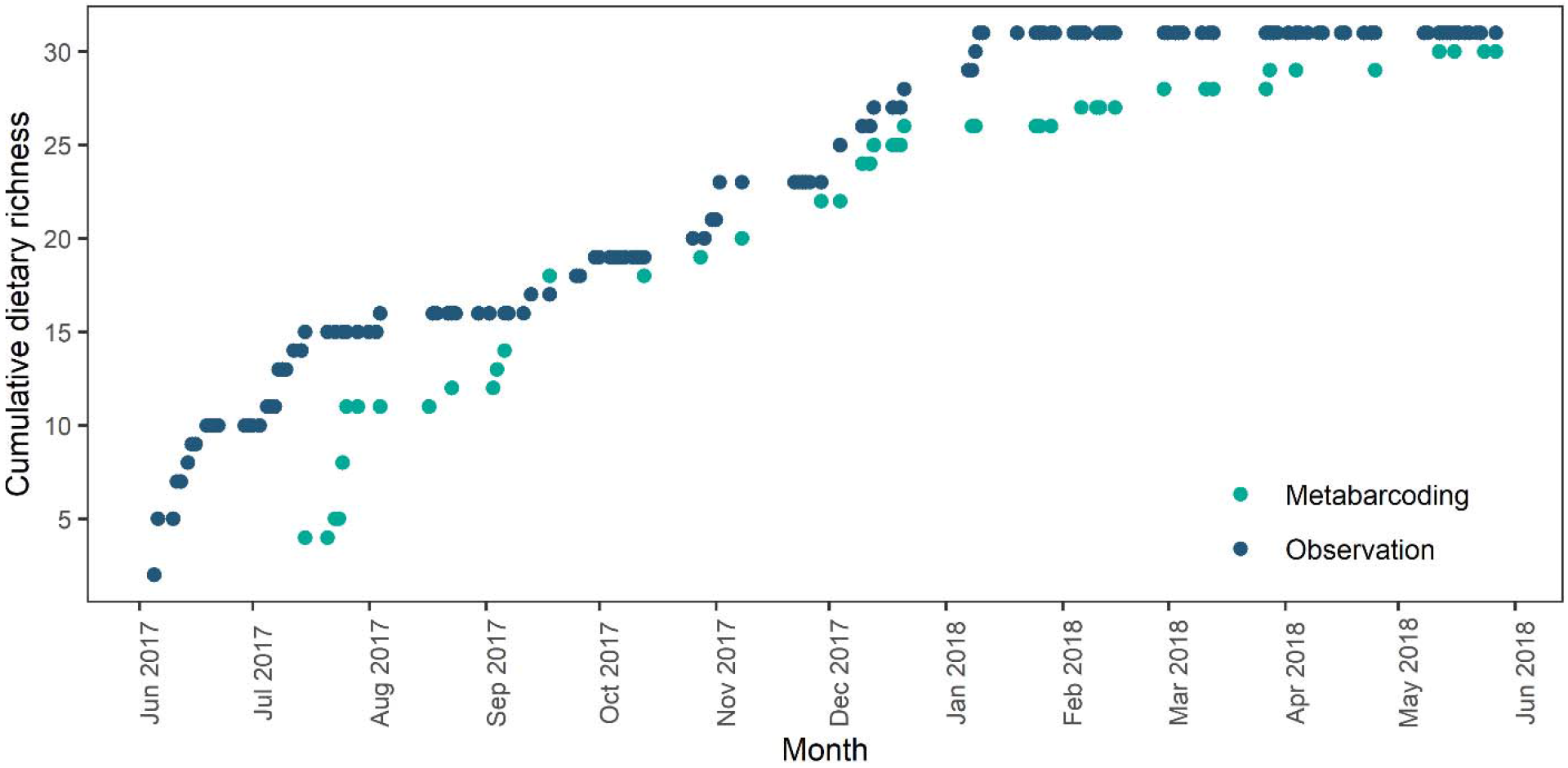
Cumulative number of resource plant taxa in the diet of reintroduced *Ara chloropterus* since the day of release based on observational (blue; *n* = 140 days) and metabarcoding (teal; *n* = 53 days) data.

**FIGURE 7.**
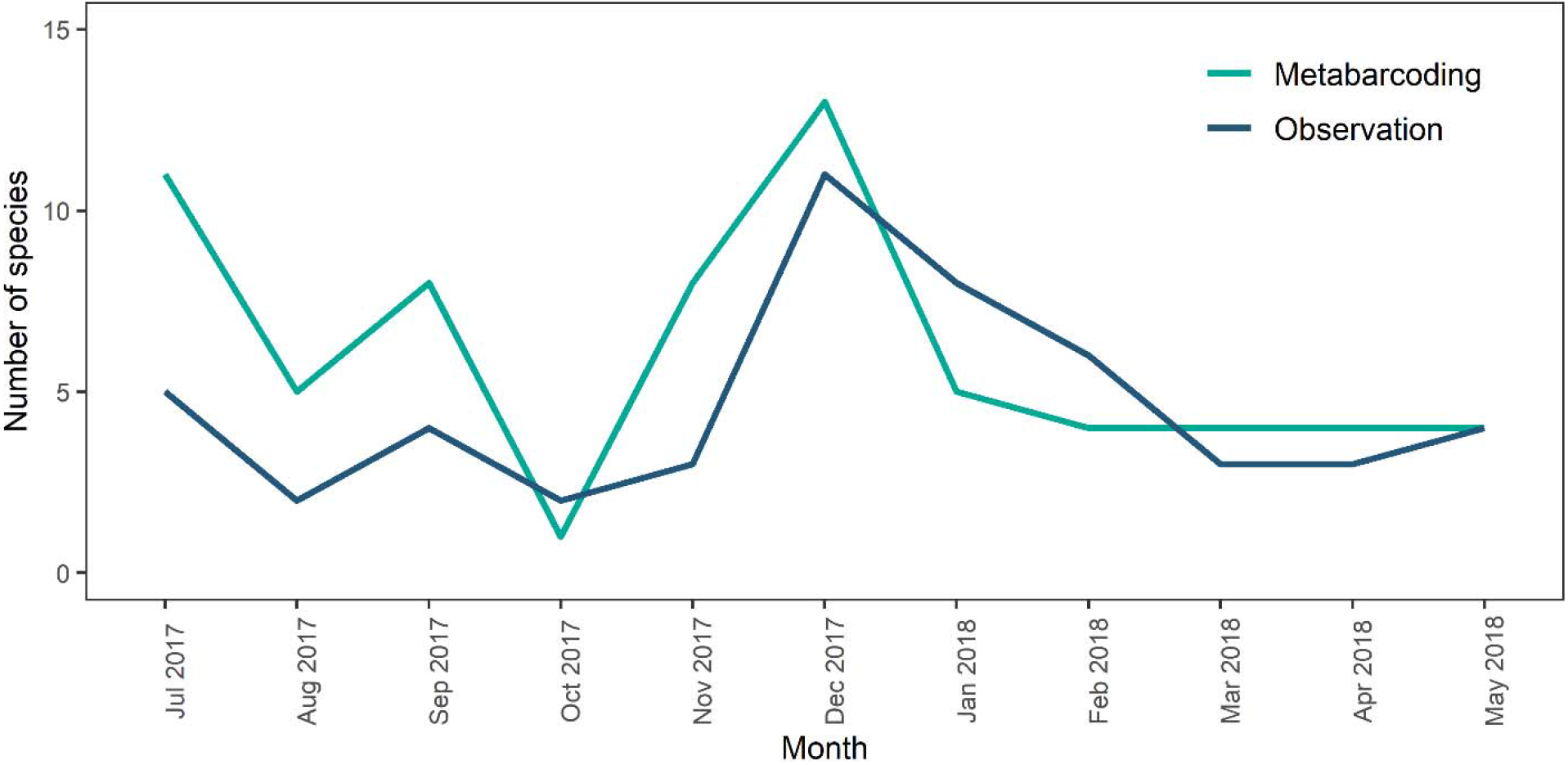
Number of resource species consumed by released Red-and-green Macaws each month, based on observational (blue; *n* = 219 feeding events) and metabarcoding (teal; *n* = 61 samples) data collected on the same date (*n* = 39 days).

**FIGURE 8.**
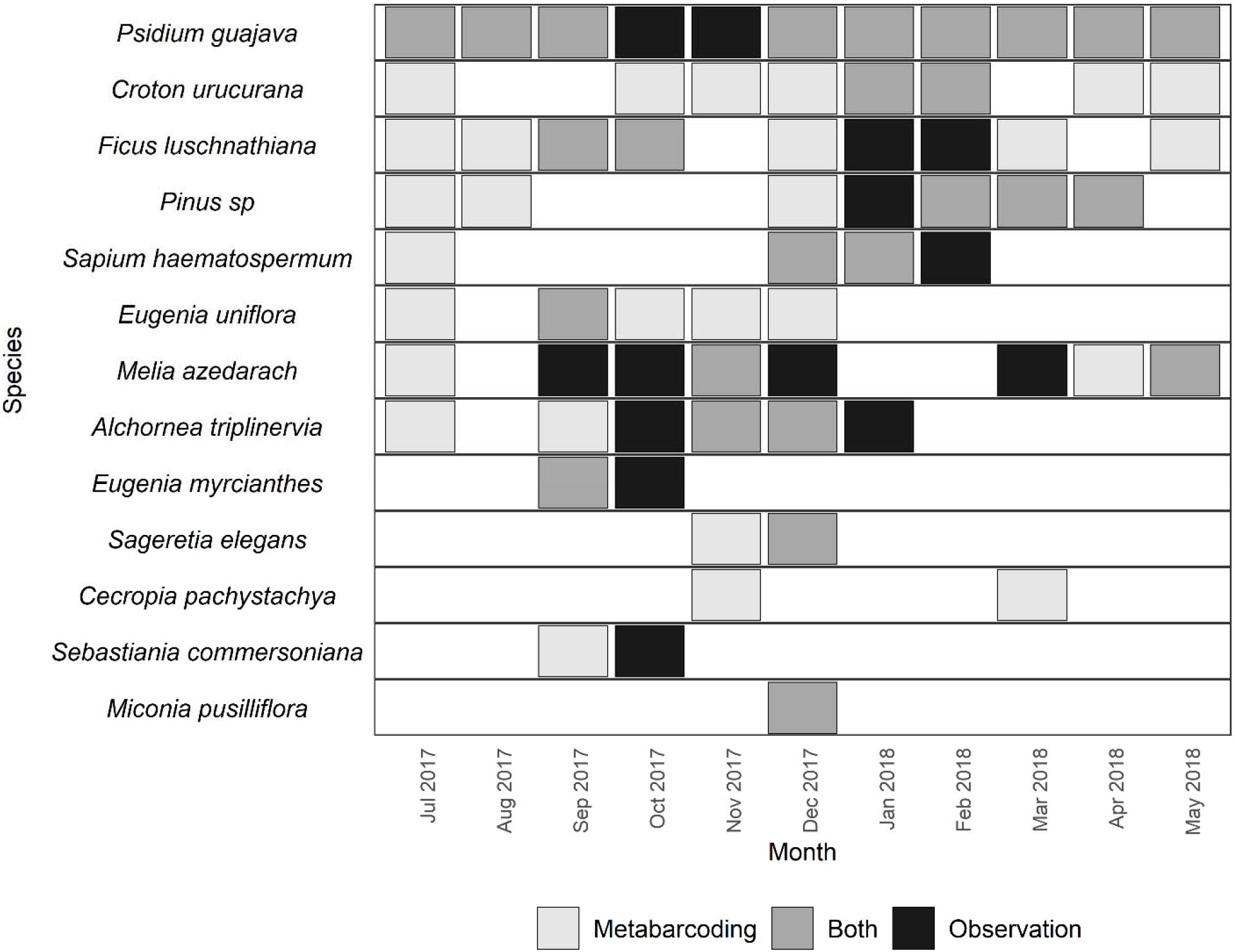
Monthly occurrence of the 13 species present in both observational (*n* = 252 feeding events) and metabarcoding datasets (*n* = 75 samples) in the diet of released Red-and-green Macaws. Only records collected within the same time frame area included. (Light grey = presence detected only by barcode, Black = presence detected only through observation, Dark grey = presence detected by both methods)

## DISCUSSION

### The Diet of Reintroduced Red-and-green Macaws

Reintroduced Red-and-green Macaws showed a good level of adjustment to life in the wild in the Iberá National Park and surrounding areas, being able to exploit a large variety of the food resources available at the site. In the one-year period studied, released macaws fed from a variety of plant species (*n* = 49) belonging to a broad phylogenetic spectrum (28 families). The observed dietary richness lies within the expected range for the species. The most exhaustive diet studies for wild Red-and-green Macaw populations to date (> 100 feeding bouts observed,>24 months of data), report a dietary breadth ranging from 10 species (Pantanal, Ferreira 2013) to 51-54 species (Amazonian rainforest, Adamek 2012; Lee et al. 2014). The overall diet composition detected in this study was similar to previous findings, with a high prevalence of detections concentrated on species from the families Fabaceae, Arecaceae and Euphorbiaceae. Additionally, 32 of the plant species eaten in the Iberá National Park belonged to the same genera as those eaten by red-and-green macaws in other regions of South America (Desenne 1994, Nycander et al. 1995, Santos 2001, Antas et al. 2002, Ragusa-Netto and Fecchio 2006, Haugaasen 2008, Adamek 2011, Scherer-Neto and Terto 2011, Ferreira 2013, Lee et al. 2014).

Macaws in this study were able to locate food sources after their release and throughout the whole year, with a monthly dietary richness that ranged from 7 to 20 species. *P. guajava* and *S. romanzoffiana* were the most frequently used plants, occurring in the diet for 12 and 11 months, respectively. The fruits and seeds of these species can be eaten by the macaws at both ripe and unripe stages and are produced year-round, making them a reliable food source. Selection for plant species with a relatively constant production of food has also been recorded for other psittacids (Bonadie and Bacon 2000, Robinet et al. 2003), with palm trees in particular being an important food source for macaws living in wetland and savanna areas (Yamashita and Machado de Barros 1997, Brightsmith and Bravo 2006, Nunes and dos Santos 2011). The strong beak of these large psittacids allows them to feed not only on the pulp but also the nuts of palm fruits (Galetti 1997), granting them access to an interior rich in lipids and proteins (Litchfield 1970, Tella et al. 2020). The use of *Psidium guajava* by macaws has seldom been reported in the literature (*Ara severa,* 2 events, Lee et al. 2014), although it is commonly used by other smaller psittacids (Paranhos et al. 2009, Silva and Melo 2013). Studies on the nutritional content of seeds of this species indicate it is a good source of proteins, as well as vitamins, and antioxidants (Uchôa-thomaz et al. 2014), while the pulp of ripe fruits has a high moisture content which can become important during the hot summer (Medina and Pagano 2003).

The composition of the diet varied along the year, responding at least in part to changes in fruit availability. For example, macaws relied on multiple species from the Myrtaceae family throughout spring (September to mid-December), switching to Euphorbiaceae during summer (December-February). This shifting between species as they become available, also known as diet switching, is the most common response to food resource fluctuation among psittacids (Renton et al. 2015), allowing them to adapt to the spatial and temporal variability of fruits, seeds and flowers. Metabarcoding results showed that some of the species were consumed outside their fruiting stage, which would confirm the ingestion of non-reproductive plant structures such as bark. The manipulation and chewing of bark was observed throughout the study, but was not considered a feeding event as it was unclear whether ingestion had occurred. The extent and function of the consumption of bark in Psittacids is yet to be determined, but it has been hypothesized to be associated with detoxification, chemical or mechanical aid in digestion, or absorption of nutrients (Warburton 2003, de Araujo and Marcondes-Machado 2011).

Two of the most frequently detected species, *Pinus elliottii* and *Melia azedarach,* were not native to the area. *Melia azedarach* has frequently been detected in the diet of psittacids living in modified landscapes (including the Red-and-green Macaw; Scherer-Neto and Terto, 2011)). Pine cones are a much less common food resource, having only been reported for one other macaw species (Silva 2018). The use of exotic plants is not uncommon in psittacids, and has allowed some species to survive and even thrive in human-dominated landscapes (Matuzak et al. 2008). They might be a useful resource when faced with a scarcity of native plants (Hamm et al. 2020), but relying on exotic species can be problematic as it can lead to conflicts with farmers (Bucher 1992) or lead to sudden population drops when plantations are harvested (Dear et al. 2010). Additionally, the presence of exotic plants is usually associated with human settlements and hence, feeding on them can increase the risk of capture by poachers, the greatest threat for psittacids (Berkunsky et al. 2017).

### Comparison Between Techniques

Dietary richness as estimated by direct observation and metabarcoding were similar (32 and 30 species respectively) though smaller than the one obtained combining the two datasets (49 species). Based on the selected observation metrics and the availability of DNA reference sequences in the study area, both techniques seem to be equally effective at describing changes in dietary richness over time, generating a similar curve of increase since time of release and similar patterns across the year.

The main difference between the results was the specific composition of the diet, with both techniques detecting at least 15 species not recorded by the other method. In the case of metabarcoding, several species, such as *S. romanzoffiana* and *I. edulis*, were expected to be present in the feces, but could not be detected albeit visual observations showed they were being consumed frequently and in large quantities by the macaws. Additionally, neither these two, nor five more species observed to be part of the diet produced an ITS2 barcode sequence during the construction of the DS-IBERAFLO database. Only ITS2 sequences for three of these species could be found at the University of Würzburg’s database. For the remaining four species, we included sequences of the respective genus and/or family in the reference database, but despite these efforts, none of the expected plants could be detected. Incomplete reference libraries always pose a challenge for taxonomic assignments of metabarcoding data, especially in tropical and subtropical habitats containing a lot of only superficially investigated biodiversity. In this study, primer specificity and plant tissue traits additionally complicate the situation. For example, ITS sequences of palm trees (Arecaceae) were not included in the original metabarcoding primer evaluation (Cheng et al. 2016) and a superficial analysis showed at least one mismatch at the forward and reverse priming site for *A. aculeata* and *S. romanzoffiana*. Furthermore, the failure to produce ITS sequences from palm trees for the custom reference database indicates a tissue-specific problem such as the presence of inhibitory substances, low DNA content, or strong cell membranes warranting additional homogenization steps during lysis. The digestive process or the potentially lower DNA quantities in the consumed palm tree nuts could further decrease the detection probability in fecal samples (King et al. 2015, Thalinger et al. 2017). All in all, the failure to detect some of the consumed species is likely a combination of these different factors. In the future, the availability of better reference databases and the development of optimized sample processing protocols will undoubtedly improve detection probabilities and the level of resolution of the results, facilitating large-scale DNA metabarcoding studies.

On the other hand, the failure to detect species by direct observation of feeding events was likely a product of observation bias. The results of an observational study can be influenced by the experience levels of observers and constrained by their ability to follow the individuals across the terrain (Ford et al. 1990, White and Garrott 2012). As a consequence, isolated food sources located in inaccessible locations will never be recorded by traditional methods, and those that are consumed only sporadically are likely to be missed.

When taking into account the sampling effort, metabarcoding showed advantages to direct observation: It had a higher detection rate than observational data, as each fecal sample was a summary of multiple feeding events. The detection rate of food items in fecal samples is influenced by the gut transition time (*i.e.* the time between its ingestion and defecation). For frugivorous birds, gut transit time has been estimated to range from a few minutes up to several hours, with longer duration in larger birds (Oehm et al. 2011, Wotton and Kelly 2012). There is currently no clear information on how long plant DNA can be detected in the feces of frugivorous birds, but studies on piscivorous birds show that prey items can be detected for up to 4 days after ingestion (Deagle et al. 2010) and that detectability is affected by meal size. When portions are small, detection rate can plummet within 24-32 hours after ingestion (Thalinger et al. 2017). Out of 48 Red-and-green Macaw fecal samples containing fruit and vegetable DNA and for which the timeframe of food availability was known, 98% contained DNA of species that had been eaten within the last 24 hours. Based on this pattern we expect each fecal sample of Red-and-green Macaws to predominantly contain remains of food ingested on either the same or the previous day, encompassing multiple feeding events. As a consequence of this high detection rate, fewer field days will be required to obtain information on dietary breadth than when working with observational data. For example, two fecal samples collected on separate days in November yielded the same number of resource species (*n* = 8) as 10 days of conducting foraging observations.

Metabarcoding also provided more detailed information on the role of particular species in the diet of the macaws, by highlighting the importance of species that were not considered significant due to an apparent low frequency of use (e.g., *Eugenia uniflora*). It also showed that the macaws were exploiting a number of wild species significantly earlier than estimated by observation (e.g., *Croton urucurana*). This information not only showed that the macaws were expanding their diet quicker than expected, but in some cases it also indicated that they were moving farther distances than detected by tracking. For example, pine trees are only located outside the boundaries of the Iberá National Park. The presence of *Pinus* sp. in fecal samples collected in July indicates that some of the macaws were feeding at forestry plantations outside of the protected area months before we were able to observe them. The downside of the technique is its inability to quantitatively compare between plant species in a fecal sample without additional tests and comparisons between food sources. Thus, estimating the relative importance of each species at any given time from DNA-based metabarcoding data is not advisable (Deagle et al. 2019, McClenaghan et al. 2019). Even if we can determine the prevalence of a species in the diet by looking at the proportion of samples in which it appears, we miss the fine-level detail that can be achieved through an observational study in which one can measure the number of items consumed, the time spent feeding on a given species, etc.

### Final Remarks

There is still no consensus on what is the optimal method to study the diet of a species, as shortcomings have been reported for all of them. Observational data can be subject to observer bias or inability to record feeding signs correctly (Shrestha and Wegge 2006, Matthews et al. 2020). Macroscopic fecal inspection is limited by the fact that mastication and digestion by the consumer can render dietary elements unidentifiable (Tutin and Fernandez 1993, Hayward 2013) while DNA-based studies are susceptible to inadequate reference databases and amplification biases (Piñol et al. 2015, Mallott et al. 2018, McClenaghan et al. 2019, Scasta et al. 2019). In the present study, we corroborate that single techniques are more limited than a combined approach in which the observation and metabarcoding of feces complement each other. By considering the results of both methods we were able to increase the number of detected food species by over 50% and better estimate the timeline of addition of novel plants in the diet after the release of the macaws. We recommend the use of a combination of observational and genetic tools in diet studies, implementing a question-oriented approach to determine the primary method of data collection. When the main focus of a study is to describe dietary composition, effort should focus on fecal sampling as metabarcoding can yield more detailed results with reduced logistical effort (assuming an adequate reference library is available). When the diet study has a behavioral component which requires information other than the identity of the consumed species (e.g., part of the plant ingested, intensity of use, etc.), then an observational approach will be needed, with fecal sampling filling the gap regarding species which are rarely eaten.

The combination of methodological approaches allowed us to establish that the Red-and-green Macaws are eating at least 49 different plant species, indicating that they have adapted well to their environment after their release and the gradual decrease in their food supplementation. This is a necessary step towards the establishment of a new population in the site. In turn, this suggests that the macaws are slowly re-gaining their dual ecological role as both regulators and disperser of seeds (Blanco et al. 2018), exhibiting a potential to affect a large diversity of local plants. On one hand, Red-and-green Macaws fed on seeds and flowers from 24 species of plants; by destructing reproductive structures macaws would be actively reducing these species reproductive output. On the other hand, macaws were seen transporting fruits for distances of up to 900m (N.L.V., personal observation), evidencing their potential role as long-distance seed dispersers. Although germination experiments would need the be conducted in order to confirm that the transported seeds are actually viable, we can expect that at least a portion of them will contribute to plant recruitment (Tella et al. 2020). In a fragmented landscape like the wetlands, where ground connectivity between forest patches is restricted by flooded terrain, seed dispersal by terrestrial vertebrates is likely limited (Nield et al. 2020). In this scenario, the presence of large bodied frugivorous birds such as the macaws can become vital to maintain gene flow between forest fragments, in particular of plant species with large seeds which cannot be transported by smaller birds (Baños-Villalba et al. 2017).

The ability of the reintroduced macaws to successfully locate and exploit food resources throughout the year, despite their captive-bred origin, can be considered as a good indicator of their acclimatization to the release site. This is an important step towards the creation of a stable, self-sustaining population of Red-and-green Macaws in the North of Argentina which can, in turn, serve as a source of individuals for the colonization of other areas from which the species has disappeared.

