## Supplementary Materials 1-2 for "Diet composition of reintroduced Red-and-Green Macaws (*Ara chloropterus*) reflects gradual adaption to life in the wild"

**SUPPLEMENTAL MATERIAL 1**

**Sample Processing**

A) DNA barcoding and reference library compilation.

The first stages of DNA Barcoding of plant tissues (DNA extraction and amplification of barcode markers) were performed at MACN following standard procedures; sequencing was done at the Centre for Biodiversity Genomics (CBG) at the University of Guelph in Canada. DNA extraction, PCR and sequencing procedures largely followed a protocol described by (Ivanova et al. 2008, Fazekas et al. 2012, Kuzmina et al. 2017). About 1–5 mg of gel-dried plant tissue was ground into fine powder using a TissueLyser II (QIAGEN) at 28 Hz for 60–90 s at room temperature, using the Axygen Mini Tube System (Axygen Scientific, Union City, California, USA) with one 3.17 mm stainless steel bead per tube. Following grinding, cells were lysed with 250 – 400 µL of 2X cetyltrimethylammonium bromide (CTAB) buffer (2% CTAB, 100mM Tris-HCl pH8.0, 20mM EDTA pH 8.0, 1.4M NaCl) incubated at 65°C for 60–90 min. After incubation, 50 µL lysate per sample was transferred into 96‐well microplates. DNA was isolated and purified using glass fiber filtration columns (Ivanova et al. 2008). PCR of an ITS fragment was performed with Phusion High‐Fidelity DNA polymerase (Fisher Scientific) using primers ITS-S2F 5’- ATGCGATACTTGGTGTGAAT-3’ (Chen et al. 2010) and ITS4 5’- TCCTCCGCTTATTGATATGC-3’ (White et al. 1990). PCR was performed following Fazekas et al protocol (Fazekas et al. 2012). Briefly, the total volume for PCR mix was 12.5 μl and included 6.25μl of 10% trehalose, 2.00 μl ddH_2_O, 1.25 μl 10X PCR buffer [200 mM Tris-HCl (pH 8.4), 500 mM KCl], 0.625 μl MgCl (50 mM), 0.125 μl of each primer, 0.062 μl dNTP (10 mM), 0.060 μl of polymerase, and 2.0 μl of DNA template. Thermal conditions were: initial denaturation at 94°C for 5 min followed by 35 cycles of denaturation for 30 s at 94°C with annealing for 30 s at 56°C and extension for 45s at 72°C; and a final extension for 10 min at 72°C. PCR products were diluted in water and sequenced at the Centre for Biodiversity Genomics, University of Guelph on an ABI 3730*xl* DNA Analyzer (Applied Biosystems) following standard procedures. Chromatograms were edited with CodonCode Aligner 3.7.1–6.0.2 (CodonCode Co.). Sequence alignments generated in MUSCLE (Edgar 2004) were used as a basis for removing primer sequences and to aid the identification of errors in sequence editing. In total, 65 of the 96 plant tissues produced ITS2 sequences corresponding to 24 different species; after filtering of contaminants and correcting for base-call errors, these were uploaded to BOLD (DS-IBERAFLO). Nine of the 33 sampled species expected to be eaten by the macaws failed to amplify. Sequences for these species, together with those of 179 other plant species occurring in the area (Arbo and Tressens 2002) were extracted from the ITS2 database hosted by the University of Würzburg (accessed 16th June 2020; Ankenbrand et al., 2015) in order to compile a custom reference database for maximum plausibility of the taxonomic assignment. In case sequences of specific species were not available, entries of the respective genus, family, tribe or order were used. The generated database was manually screened for misidentified sequences, which were subsequently removed.

B) **Metabarcoding of fecal samples.**

Metabarcoding of the fecal samples was carried out at the CBG. Feces were processed in a laboratory dedicated to the handling of low-quality DNA samples with separate rooms for DNA extraction, PCR preparation and post-PCR processing. DNA-free gloves were worn at all times and all surfaces were cleaned with 0.6% sodium hypochlorite followed by 70% Ethanol prior and after use. Additionally, fecal samples were only opened under PCR hoods, which were UV-ed for 15 min prior to any processing step. After removing the ethanol from the falcon tubes, samples were dried in a fume-hood. One empty reaction tube with an open lid was placed adjacent to the fecal samples. This hood-control was processed together with the other samples to detect possible cross-contamination during the drying process. Subsequently, samples were transferred to new reaction tubes for further processing. For lysis, a 40:1 mix of insect lysis buffer ( 700mM GuSCN, 30mM EDTA pH 8.0, 30mM Tris-HCl pH 8.0, 0.5% Tritón X-100, 5% Tween-20) and Proteinase K (20 mg/ml) was used (Ivanova et al. 2008). Depending on the size of the fecal sample, 300µl - 6ml of the mix was added to cover the fecal sample entirely. Samples were incubated overnight at 56°C on an orbital shaker. For DNA extraction with the DNeasy kit (QIAGEN), we used 200µl of lysate and followed manufacturer’s instructions, except for an additional centrifugation step removing AW2 Buffer from the membrane and for elution with Buffer AE in two steps of 100µl each. An extraction control only containing lysis buffer was included in the extraction process and successful extraction was confirmed by measuring the total DNA concentration with the Qubit dsDNA HS Assay Kit (Thermo Fisher Scientific, MA, USA).

All DNA extracts were subjected to a metabarcoding approach using two consecutive PCRs and fusion primers (Elbrecht and Steinke 2019), for which cycling conditions were optimized for maximum yield of target length fragments, while minimizing the occurrence of non-target bands. We included fourteen negative controls in both PCRs. The first round of PCR employed the primers ITS-u3 5’-CAWCGATGAAGAACGYAGC-3’ and ITS-u4 5’-RGTTTCTTTTCCTCCGCTTA-3’ (Cheng et al. 2016). Each 25 µl reaction contained 1.25 µl of 2× Multiplex Master Mix Plus (QIAGEN, Hilden, Germany), 1.25 µl of each primer (10 µM), 4 µl DNA extract, and molecular grade water. Thermocycling conditions were 5 min of initial denaturation at 95°C, followed by 24 cycles of 95°C for 30 s, 50°C for 30 s, and 72°C for 50 s, and final extension at 68°C for 10 min. Two µL of PCR product were used as template for the second PCR, where Illumina sequencing adapters were added using individually tagged fusion primers (Elbrecht and Steinke, 2019; Supplementary Material Table S1). Every 25 µl reaction contained 12.5 µl of 2× Multiplex Master Mix Plus (QIAGEN, Hilden, Germany), 1.25 µl each fusion primer (10 µM), 2 µl PCR template, and molecular grade water. Thermocycling conditions were identical to round one. Successful amplification was confirmed by visualizing amplicons on 1.5% agarose gels and noting target fluorescence (none, weak, strong).

PCR products were purified and normalized using SequalPrep Normalization Plates (Thermo Fisher Scientific, MA, USA), (Harris et al. 2010) according to the manufacturer’s instructions. Fifteen µL of each normalized sample were pooled, and the final library cleaned up using left-sided size selection with 0.76x SPRIselect (Beckman Coulter, CA, USA). The average DNA concentration of the final library was 6.96 ng/µl (as measured in triplicate with Qubit Fluorometric Quantification) and the presence of target length fragments was confirmed on a 1.5% agarose gel. Sequencing was carried out by the Advanced Analysis Centre at the University of Guelph using a 600 cycle Illumina MiSeq Reagent Kit v3 and 5% PhiX spike in. Altogether, 112 samples and negative controls were sequenced simultaneously leading to an average expected sequencing depth of approx. 220,000 reads per sample. Sequencing results were uploaded to the Sequence Read Archive (SRA, Genbank, accession: PRJNA695029).

C) **Bioinformatic analyses.**

Resulting sequence data were processed using the JAMP pipeline v0.67 (github.com/VascoElbrecht/JAMP). Sequences were demultiplexed, paired-end reads merged using Usearch v11.0.667 with fastq_pctid=75 (Edgar 2010), reads outside a 100 bp to 430 bp range were discarded and primer sequences trimmed by using Cutadapt v1.18 with default settings (Martin 2011). Sequences with poor quality were removed using an expected error value of 1.5 (Edgar and Flyvbjerg 2015) as implemented in Usearch. All sequences with less than five reads were removed during the denoising process.

The obtained haplotypes were mapped against the custom sequence database: First, only 100% matches were identified, the remaining reads were subsequently mapped using identity thresholds of 98% 95%, 90% and 80%. The permitted taxonomic resolution was determined by two factors: Based on the quality of the reference database and based on the percent identity score. In case reference sequences were only available at the genus level, but not for the plant species occurring in the study area, the genus level was the highest taxonomic resolution possible, independent of the identity score. For identity scores of 80-90%, 91-95%, 96-97% and 98-100% the taxonomy was determined on the order, family, genus and species level, respectively, independent of the local species composition.

The remaining haplotypes without matches in the custom reference database were blasted and results filtered using the first 500 hits with a bit-score above 155, of which only the hits with a bit-score above 98% of the bit-score of the top hit were kept. Based on the percent-identity, the taxonomic resolution of the remaining hits was determined with 80-84%, 85-89%, 90-94%, 95-96%, 97-98% and 99-100% for phylum, class, order, family, genus and species level. In comparison to the custom reference sequence database, a stricter taxonomic cut-off was used. Using a lowest common ancestor approach the taxonomic level for which all resulting hits shared their taxonomic assignment was determined for each haplotype. Fungi and Bacteria were not further identified, but haplotypes mapping to Viridiplantae were combined with the results of mapping against the custom sequence database.

The detected taxa were classified into five categories: Resource = wild local plants known to be a food item or considered likely to be so based on its characteristics (fruit-producing tree or vine); Provided = commercial fruits or vegetables included in the daily food supplements; Provided/Resource = level of resolution did not allow to exclude either option; Contamination = unlikely to have been eaten by the macaws (included algae and herbaceous or aquatic plants) and Ambiguous = could be either of the previous categories. Only taxa classified as Resource were included in the dietary analysis.

**SUPPLEMENTAL MATERIAL 2**

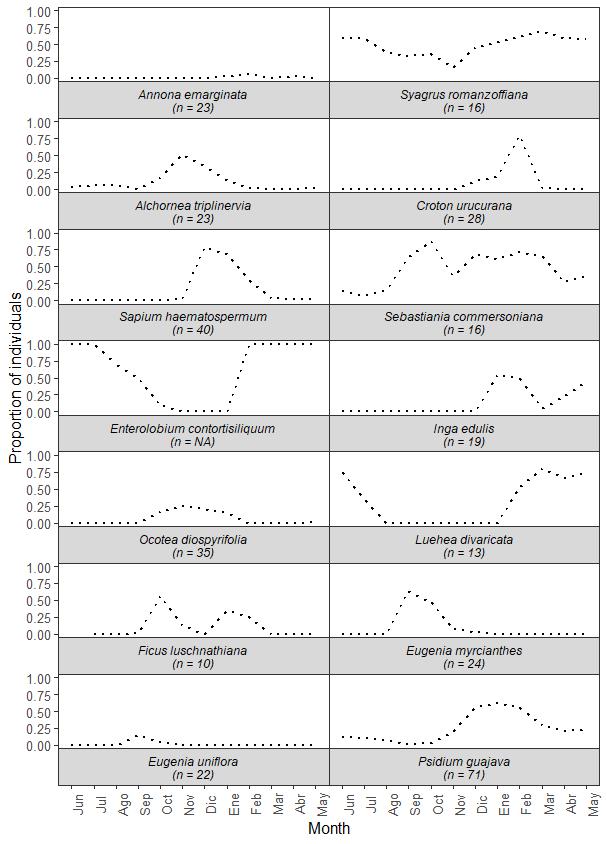

**FIGURE S2.** Availability patterns for 14 species observed being consumed by released *Ara chloropterus*, estimated as the proportion of sampled individuals observed at a pheno-phase edible by the macaws on a given month. The value *n* indicates de number of individual trees visited monthly, except for *Enterolobium contortisiliquum* whose phenology is based on a combination of non-systematic observations and bibliographical data (Crechi et al. 2010).

**TABLE S2.** Results of Spearman rank correlation tests evaluating the relationship between the monthly number of feeding events and the proportion of trees of a given species producing fruits edible by *Ara chloropterus.* Rho values measure the strength of the association, with values close to 1 or -1 indicating a strong positive or negative correlation respectively.

| **Species** | **Rho** | **p** |
| --- | --- | --- |
| Annona emarginata | 0.47 | 0.13 |
| Syagrus romanzoffiana | -0.07 | 0.83 |
| Alchornea triplinervia | 0.83 | **<0.001** |
| Croton urucurana | 0.66 | **0.02** |
| Sapium haematospermum | 0.78 | **<0.001** |
| Sebastiania commersoniana | 0.48 | 0.11 |
| Enterolobium contortisiliquum | 0.56 | 0.06 |
| Inga edulis | 0.71 | **0.01** |
| Ocotea diospyrifolia | 0.75 | **0.01** |
| Luehea divaricata | 0.09 | 0.77 |
| Ficus luschnathiana | 0.74 | **0.01** |
| Eugenia myrcianthes | 0.77 | **<0.001** |
| Eugenia uniflora | 0.57 | 0.05 |
| Psidium guajava | 0.13 | 0.69 |
